# Crystal Structure of Caskin1/CASK complex reveals the molecular basis of the binding specificity of CASK_CAMK domain and its binding partners

**DOI:** 10.1101/2022.01.16.476467

**Authors:** Yue Wang, Qiangou Chen, Riting Jiang, Xiaoyang Ye, Jun Wan, Jianchao Li, Wei Liu

**Affiliations:** Shenzhen Key Laboratory for Neuronal Structural Biology, Biomedical Research Institute, Shenzhen Peking University-The Hong Kong University of Science and Technology Medical Center, Shenzhen 518036, China; Division of Cell, Developmental and Integrative Biology, School of Medicine, South China University of Technology, Guangzhou, China; Department of Otorhinolaryngology, Guangzhou First People’s Hospital, School of Medicine, South China University of Technology, Guangzhou, China

**Keywords:** CASK, CAMK, Caskin1, interaction, crystallization

## Abstract

CASK is a unique scaffold protein in the synapse system. It links numerous proteins to the pre- or post-synaptic region and is critical to the homeostasis of synaptic vesicles. The N-terminus of CASK is a calcium/calmodulin-dependent protein kinase (CAMK) domain, which has diverse functions and interacts with downstream proteins to form a scaffold platform. Caskin1 is one of the brain-specific adaptor proteins of CASK. Previous studies showed that CASK_CAMK domain interacts with Caskin1 CID domain with relatively low affinity. In this study, we re-visit this interaction by remapping the interaction boundary and solving their complex structure. Based on the structure, we systematically compared the interactions between CASK_CAMK and other binding partners. Our results showed that CAMK domain occupies the CID peptide by using its C-lobe groove (between the α1 and α2) and there is a highly conserved signature motif (ζ-x-ψ-W-ψ-x-R) in the CID domain, where ζ is acidic side chain containing residues, x is any amino acid residue, ψ is hydrophobic residues, W is for tryptophan, and R is arginine. These findings allowed us to identify several new potential cytoplasmic binding partners for CASK_CAMK.

## Introduction

CASK (Calcium/calmodulin-dependent serine protein kinase) is a vital unique scaffold protein in the synapse, which was first identified as neurexin’s adaptor protein [1–3]. In the brain, CASK is localized in pre- and post-synaptic densities [4]. As a scaffold protein, CASK participated in multiple interactions by utilizing its different domains [5–8]. The N-terminus of CASK contains a calcium/calmodulin-dependent protein kinases (CAMK) domain, followed by a tandem Lin2/Lin7 domain; its C-terminal half is composed of a PDZ domain (post-synaptic density protein (PSD95), Drosophila disc large tumor suppressor (Dlg1), and zonula occludens-1 protein (zo-1)), SH3 domain and guanylate kinase domain [9, 10]. According to its domain organization, it was identified as a CAM kinase and a scaffold domain as well, suggesting multiple functions that may be involved in different cell biological events.

In C.elegans, CASK was identified as the product of lin-2 [4], connected with Lin-10 (aka Mint or X11) and Lin-7 (aka MALS or Veli) [11], form a critical platform for vulval induction in organizing synapses. In Drosophila (where it is called CamGUK), its mutation causes synaptic defects and behavioral dysfunctions [12]. Consistent with this notion, CASK binds to neurexins, SynCAMs, and SAP97 in mammalian cells [13–15]. Knocked out CASK in mice is lethal. Even reducing CASK expression will impair synaptic function [16]. In humans, CASK-related disorders are not limited by neurodevelopmental syndromes, such as X-lined mental retardation, microcephaly, and autism [17–19]. Cooperating with other scaffold proteins, like discs-large homolog 1 (DLG1), CASK also plays a critical role in kidney development [20].

Caskin1 was first identified as a brain-specific adaptor protein for CASK [21]. It has a homolog protein, Caskin2. The two proteins have similar domain organization. Their N-terminus contain Ankyrin Repeats, followed by an SH3 domain [22] for lipid binding, a CASK interaction domain (CID), and a SAM domain tandem [23]. In comparison, the C-terminus is the Proline-rich unstructured region.

In this study, we first re-mapped the interaction between CASK_CAMK and Caskin1_CID and solved the high-resolution crystal structure of CASK_CAMK/Caskin1_CID complex. We demonstrated that the interaction depends on the C-lobe groove (between the α1 and α2) of CASK_CAMK domain and a highly conserved ζ-x-ψ-W-ψ-x-R motif in CID, where ζ is acidic side chain containing residues, x is any amino acid residue, ψ is hydrophobic residues, W is for tryptophan, and R is arginine. Using this model, we searched for new binding partners among all human proteins. We found several potential candidates and confirmed the interactions *in vitro*. CIC and LRRK2 are involved in brain development and highly expressed in brain. Given this background and the interaction we verified, the two proteins may interact with scaffold protein CASK and function in brain development.

## Material and Methods

### 1. Materials

All DNA restriction enzymes, DNA clean up and gel extraction, and plasmid extraction kits were purchased from New England Biolabs. DTT, Tris and EDTA were obtained from Sigma. Other chemicals of analytical reagent grade were bought from Sangon Biotech.

### 2. Plasmid

The cDNAs-encoding CASK (NM_009806.3), Caskin1(NM_027937.2), Mint1(NM_177034.3), Tiam1(NM_009384.3), LRRK1 (NM_146191.3), and RP1L1 (NM_146246.3) from Mus musculus were amplified from a mouse cDNA library (Gift from Dr. Jun Wan’s Lab). Primers used in this study were listed in Table S1. The PCR-amplified productions were subcloned into a modified pET28a or pET32a vectors [24].

### 3 Protein expression and purification

The plasmids encoding the recombinant CASK_CAMK domain protein were transformed into *Escherichia coli* BL21 (DE3). Single colony was inoculated to LB media containing 50μg/ml Ampicillin. The cells were induced with 0.1mM IPTG at 16 °C for 20 h when O.D.600 reached to 0.6~0.8. After induction, the cells were harvested by centrifugation at 5000g for 15 min, lysed with a high-pressure homogenizer (UH06, union-biotech, China) and finally centrifuged at 20,000g for 20 min to collect the supernatant. The His-tagged fusion proteins were purified by Ni^2+^-Sepharose 6 fast flow (GE Healthcare, USA) affinity chromatography followed by size-exclusion chromatography.

### 4 SDS-PAGE electrophoresis

SDS–PAGE was performed using a 5% stacking gel and a 12% separating gel in Tris–glycine buffer. Gels were run in a vertical electrophoresis Mini-PROTEAN Tetra Cell apparatus (Bio-rad, Austria) and stained with Coomassie blue. The protein mass standard used was the PageRuler Prestained Ladder (Fermentas, Austria).

### 5. Size exclusion chromatography and Multi-angle Laser Light Scattering (SEC-MALS)

Size exclusion chromatography was performed on AKTA system using Superose 12 10/300 GL column, coupled to a Dawn Heleos-II light scattering detector (Wyatt Technology Corporation, CA) and an Optilab-Rex refractive index monitor (Wyatt Technology Corporation). Molecular mass calculations were performed using ASTRA software (Wyatt Technology Corporation). Protein samples were injected into the column pre-equilibrated with 50 mM Tris, pH 7.5, 100 mM NaCl, 1 mM DTT, and 1 mM EDTA.

### 6. Isothermal titration calorimetry assay

Isothermal titration calorimetry experiments were carried out on the MicroCal ITC200 calorimeter (Malvern, UK) at 25°C. The concentration of the injected samples in the syringe was 500 μM, and the concentration of the samples in the cell was fixed at 50 μM. The sample in the syringe was sequentially injected into the sample cell with a time interval of 150 s (0.5 μL for the first injection and 2 μL each for the following 19 injections). The titration data were analyzed by the Origin 7.0 software and fitted with the one-site binding model.

### 7. Cellular colocalization and co-immunoprecipitation

HEK293T cells were cultured in Dulbecco’s Modified Eagle Medium (DMEM) supplemented with 10% fetal bovine serum (FBS), and 1% of penicillin-streptomycin at 37°C with 5% CO2. HEK293T cells were transfected with GFP-tagged full-length wild type CASK, with or without RFP-tagged full-length Caskin1 using ViaFect Transfection Reagent (Promega, Madison, WI) as per manufacturer’s protocol. These cells were not individually authenticated and not found to be on the list of commonly misidentified cell lines (International Cell Line Authentication Committee). Cells were tested negative for mycoplasma contamination by cytoplasmic DAPI staining.

The cells were then fixed for 15 min with 4% paraformaldehyde in PBS. Protein colocalizations were analyzed with a Zeiss Confocal microscope (LSM710) equipped with a digital camera with a 100× objective lens. The cell areas were measured using the Fiji [25].

HEK293T cell lysates expressing the Flag-tagged full length Caskin1, and HA-tagged full length CASK proteins were incubated with 1μg anti-Flag M2 antibody (Sigma, F3165) at 4°C over night. The mixture was centrifuged in 14,000 rpm for 10mins at 4°C, then loaded the supernatant onto 20 μL Protein A beads (GE Healthcare) for 2h in cold room. After washing twice with PBS plus 0.1% Triton, the proteins captured by the beads were eluted by boiling with SDS-PAGE loading buffer, resolved by SDS-PAGE and detected by an anti-Flag and anti HA antibodies (CST, 3724) using western blotting.

### 8. Protein crystallography

The CASK_CAMK complex were concentrated to 20 mg/mL mixed with Caskin1_CID 371-381 peptide on the stoichiometry of 1:1.5 for crystallization. Crystals were obtained at 16°C by the sitting-drop vapor diffusion against 80 mL well solution using 48-well format crystallization plates. The complex crystals were grown in a buffer containing 2.0 M Ammonium citrate tribasic pH 7.0, 0.1 M BIS-TRIS propane pH 7.0. Crystals were soaked in the crystallization solution with saturated concentration of Ammonium citrate for cryo-protection. Diffraction data were collected at the Shanghai Synchrotron Radiation Facility BL17U1 at 100 K. Data were processed and scaled using HKL2000 [26].

Structure was solved by molecular replacement with the CASK_CAMK domain (PDB: 3TAC) as the searching model using PHASER [27]. Further manual model buildings and refinements were completed iteratively using Coot [28] and PHENIX [29] or Refmac5 [30]. The final model was validated by MolProbity [31]. The final refinement statistics are summarized in Table 1. The structure figures were prepared by ChimeraX [32].

## Results

### 1. Mapping the interaction between CASK_CAMK and Caskin_CID

The schematic domain organization of CASK and Caskin1 is shown in Figure 1.A. Caskin1, which has been reported as a brain-specific binding partner for CASK [21], the protein has markable structure features. According to the predicted domain organization and conserved sequence analysis, Caskin1 could be divided into two parts. The N-terminus is the well-folded Ankyrin-Repeats domain, followed by an SH3 domain, CASK-interaction-domain (CID), and a tandem SAM domain; the C-terminus is a disordered non-folding proline-rich region.

**Figure 1.**
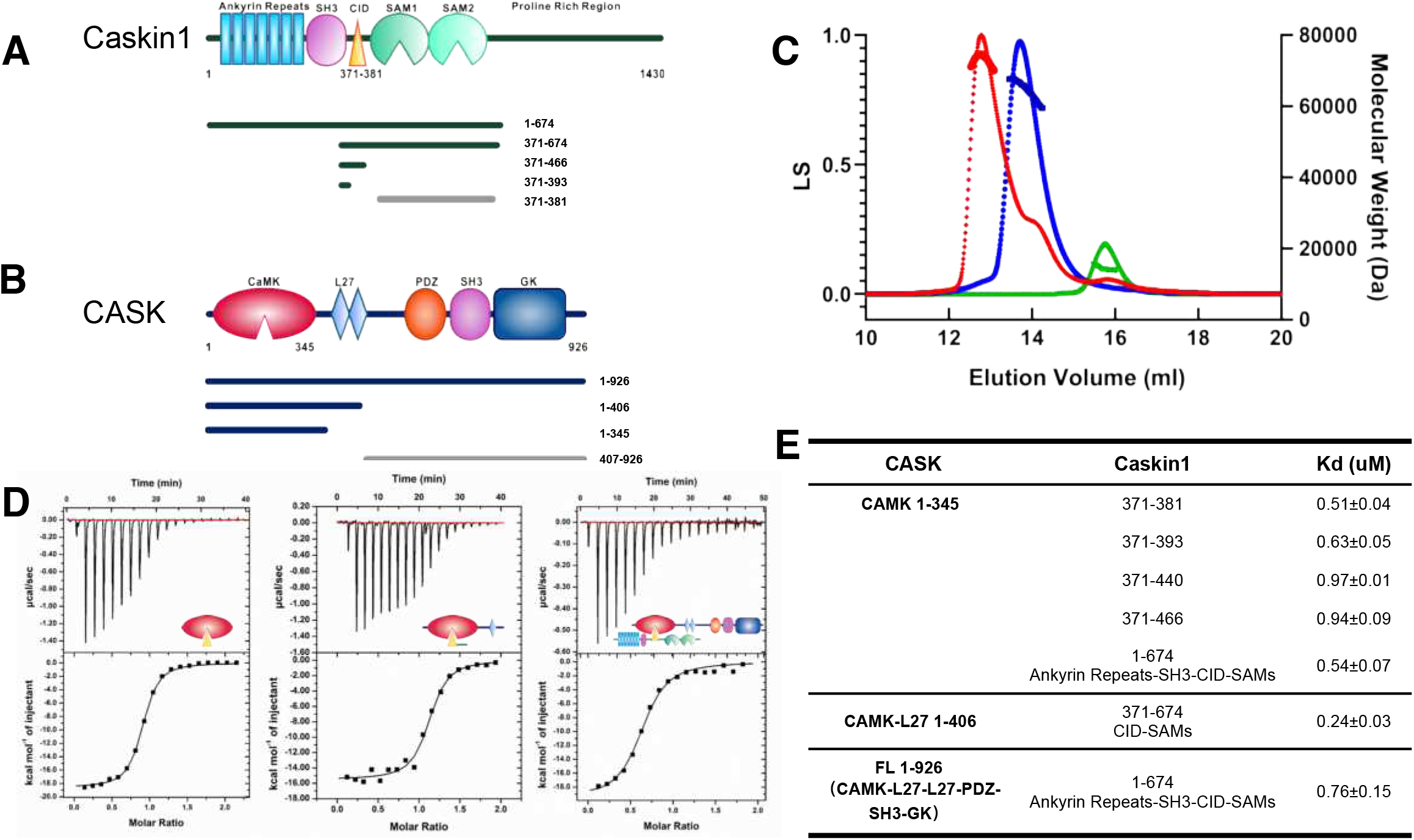
Caskin1_CID interacts with the CASK_CAMK domain. (A-B) Schematic diagrams represent the domain organizations of Caskin1 and CASK. Underline represented the variant fragments we adapted.(C) SEC-MALS assay presented Caskin1 (residues 371-466 with Trx-his tag, and the molecular weight is determined to be ~13.20kD) and CASK_CAMK (residues 1-345 with his tag, is ~59.90kD) formed a stable complex (~65.83kD) with stoichiometry about 1:1. (D) ITC-based measurements of the interaction between Caskin1 and CASK. (E) Summary of the dissociation constants between Caskin1 and CASK.

We first confirmed the interaction between Caskin1 and CASK. The recombinant CASK full-length protein and the N-terminus of Caskin1 were expressed in *E. coli* BL21(DE3) system. According to previous research, the Caskin1 binds to the CASK_CAMK domain by using a linear peptide [33]. Our ITC (Isothermal Titration Calorimetry) result revealed that the reported Caskin1 peptide bound to CASK_CAMK with a dissociation constant (Kd) of ~6.94 μM at a 1:1 stoichiometry, which is consistent with the previous study.

From sequence analysis of Caskin1, we noticed that the short peptide might not fully cover the interaction region (Figure S1). The other highly conserved amino acids may also be involved in the interaction with CASK. To test this hypothesis, we expressed a series of Caskin1 proteins and evaluated the interaction with CASK_CAMK. Compared with the previous reports, the extended CID protein (residues 371-466) binds to CASK_CAMK domain (residues 1-345) with a higher binding affinity (0.94±0.09 μM). Full-length CASK protein bound to Caskin1_N-terminal (residues 1-674) with the dissociation constant of ~0.76 μM, similar with the CAMK domain alone (Figure 1.D). In the size-exclusion chromatography coupled with multi-angle light scattering (SEC-MALS) experiments, both CASK_CAMK (residues 1-345 with his tag) and Caskin1 (residues 371-466 with his tag) proteins showed sharp and symmetrical peaks and their molecular weights were highly homogenous (Figure 1.B).Therefore the purified CASK and Caskin1 proteins were suitable for further biochemical characterization experiments. In addition, the Caskin1/CASK complex formed a stable complex with 1 to 1 stichometry (Figure 1.C). Further truncation of Caskin1 revealed that Caskin1 371-381 peptide contained the complete CASK binding components. Therefore, we concluded that Caskin1 CID (residues 371-381) and CASK_CAMK (residues 1-345) are necessary and sufficient to form the CASK/Caskin1 protein complex (Figure 1.E).

Since Caskin1 was identified as the second CASK_CAMK binding protein in the brain besides Mint1[21], we also validated the Caskin1/CASK interaction *in vitro* by colocalization analysis and co-immunoprecipitation assay (Figure S2.A). In HEK293T cell, we expressed the Caskin1 full-length protein with RFP-tag and GFP-tagged CASK. Caskin1 is well co-localized with CASK under confocal microscope observation, providing the *in vitro* evidence that Caskin1 binds to CASK (Figure S2.B).

### 2. Crystal structure of CASK_CAMK and Caskin1_CID complex

To further illuminate the interaction in detail and find the key residues for CASK/Caskin complex, we crystallized the CASK_CAMK and Caskin1_CID 371-381 peptide. The 2.5Å crystal structure allowed us to identify the interactive features of the CASK/Caskin1 complex (Table 1).

The overall structure of the CASK/Caskin1 complex adopts a typical CAMK domain/CID binding mode, in which the CID peptide is buried into the CAMK C-lobe (Figure 2.A). The structures determined in this study were also consistent with the previous docking prediction [33] and the CASK/Mint1 studies. By superimposing the three structures, we found that they all adopted a similar binding model (Figure S3) [34, 35].

**Figure 2.**
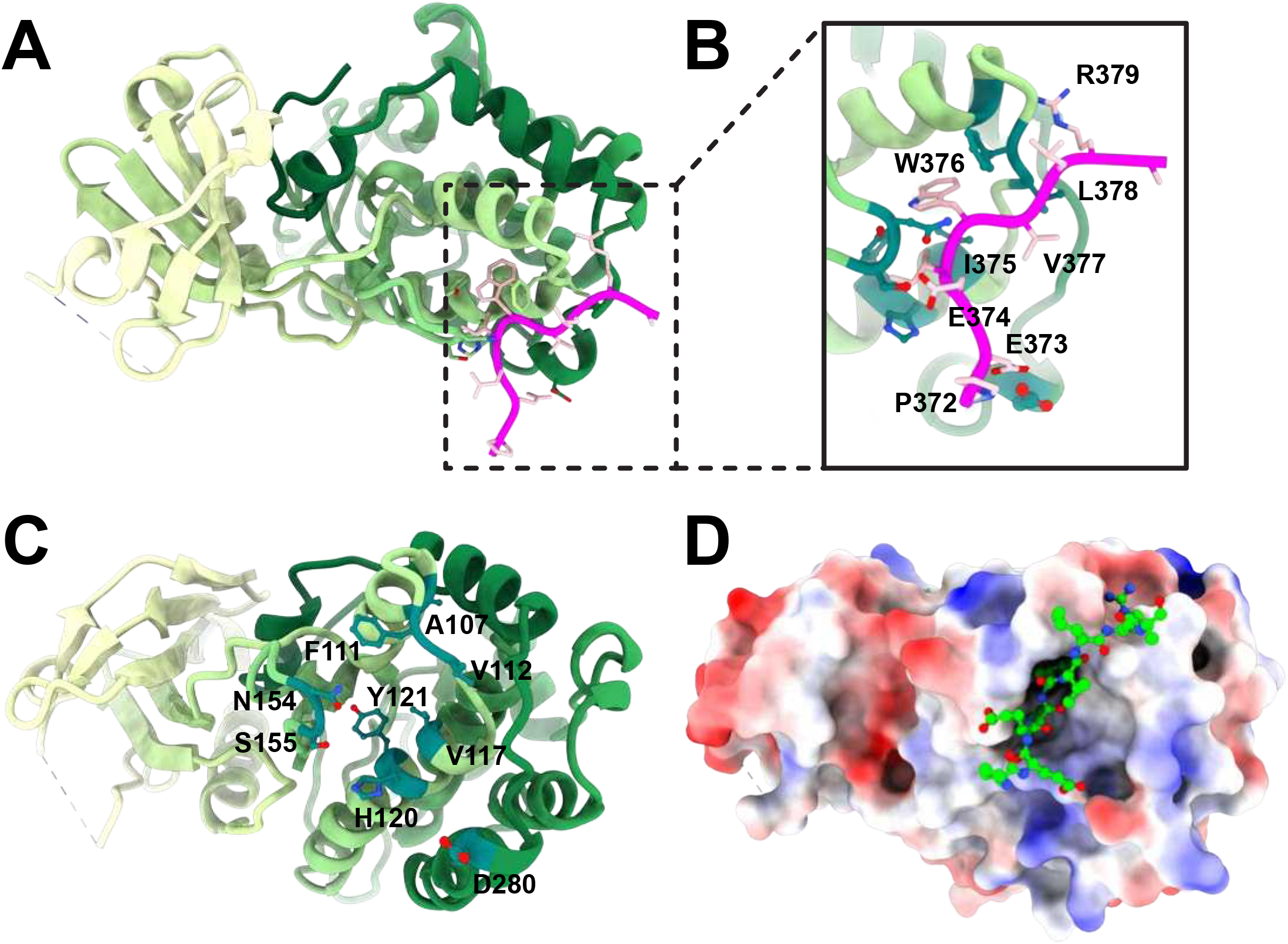
The detailed interactions governing the formation of Caskin1_CID and CASK_CAMK coomplex. (A) Ribbon representations of the Caskin1_CID(residues 371-381) and CASK_CAMK (residues 1-345). (B-C) Details of the interfaces in the Caskin1_CID (C) and CASK_CAMK (C). (D) Caskin1 is buried into the hydrophobic groove of Cask. Molecular surface representation showing the electrostatic surface potential of CASK_CAMK. Caskin1_CID shows in ball-sticks.

In the high-resolution structure, we revealed that Caskin1 W376 and I375, combined with CASK V117, H120, and Y121 to form a hydrophobic core (Figure 2.B and C). Replaced any one of these residues, the complex will be damaged. Caskin1 W376 is strictly conserved in all CID-containing proteins, and it was deeply buried into CASK hydrophobic pocket (Figure 2.D). ITC-based measurements further demonstrated that the substitution of W376 will totally abolish the interaction.

### 3. The key residues in the CASK_CAMK/CID interface

CASK_CAMK domain has been reported to have various binding partners, such as Mint1, Caskin1, Tiam1, and LiprinA2. Here, we compared the CID peptides sequence among these targets (Figure 3.A). Based on ITC measurements, the Caskin1 CID has the strongest binding affinity. While Tiam1 and Mint1 CID interact with CASK_CAMK domain with lower binding affinities (Figure 3.B).

**Figure 3.**
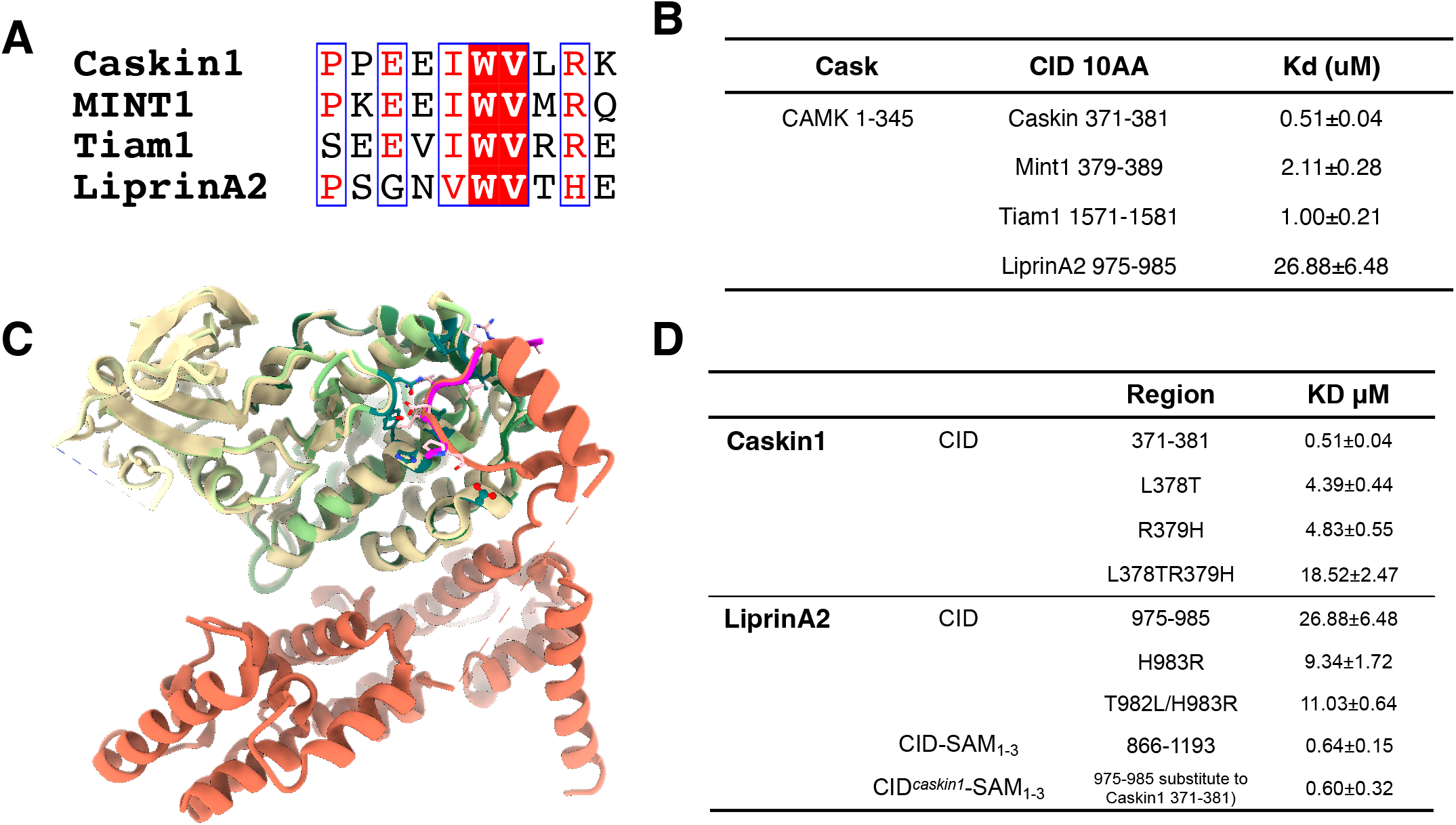
Comparision of four CID-containing proteins and the following mutagenesis study. (A) Sequence alignments of four CID-containing proteins. (B) Summary of ITC-derived dislocation constants between CASK_CAMK and the four CID proteins.(C) Superposition of CASK_CAMK/LiprinA2_SAMs with CASK_CAMK /Caskin1_CID. (D) Mutations of Caskin1 CID and LiprinA2 CID reveal the key residues.

Superimposition of the Caskin1/CASK structure with the LiprinA2/CASK structure reveals that the CID insertion between SAM1 and SAM2 of LiprinA2 is critical (Figure 3.C). We then ask whether we can further increase the LirpinA2 CID binding capacity by introducing the critical amino acids of the Caskin1 into LiprinA2 CID. L378^Caskin1^ and F111^CASK^ provide the hydrophobic interaction between CASK/Caskin1. If we mutated the L378^Caskin1^ to hydrophilic T^LiprinA2^, the interaction would decrease by about 9-folds. R379 ^Caskin1^ positive charged side chain stacked with H278 ^CASK^ imidazole side chain. Mutating the R379 ^Caskin1^ to H or L ^LiprinA2^ will slightly reduce the binding affinity. Double mutation L378T/R379H will dramatically decrease the binding by ~40-folds, meaning that these two amino acids in Caskin1 are important for CASK binding. At the same time, gain-of-function experiments showed that LiprinA2 G977E^liprinA2^ and H983R^liprinA2^ mutations led to ~9 folds increase in the binding affinity to CASK. However, the substitution of LiprinA2 CID by Caskin1 CID in the LiprinA2 protein did not further increase the interaction, indicating that the CID and SAMs binding interfaces are independent of each other (Figure 3D).

### 4. Identification of other potential candidates for CASK_CAMK binding

CASK plays not only vital roles in neuron development and synapse vesicle transport but also has essential functions in other organs, especially the vesicle transport function for insulin secrete has been proved associated with CASK [34, 36].

Based on the crystal structure of CASK/Caskin1 complex and CID sequence analysis of known binding partners, we proposed a binding motif in CID for CID/CASK interaction: ζ-x-ψ-W-ψ-x-R, where ζ is acidic side chain containing residues, x is any amino acid residue, ψ is hydrophobic residues, W is for tryptophan, and R is arginine. The ζ-x-ψ-W-ψ-x-R peptide-containing protein should be potential binding partners of CASK_CAMK.

Using this model, we searched the protein database and found several potential candidates that may interact with CASK_CAMK in the protein database (Figure 4.A), including capicua homolog (CIC), Leucine-rich repeat serine/threonine-protein kinase1 (LRRK1), FAM200A and Rp1l1. We purified the CIC and LRRK1 CID protein with Trx-His-tag and tested the interaction with CASK_CAMK. Indeed, CIC and LRRK1 have a mild interaction with CASK_CAMK (Figure 4B). FAM200A has an unknown function membrane protein and Rp1l1 plays a role in the organization of the outer segment of rod and cone photoreceptors. We purified these CID-containing proteins and measured their individual bindings to CASK_CAMK by ITC-based assays. The results are summarized in (Figure 4.B). FAM200A and Rp1l1 have a suitable hydrophobic sequence. However, the C-terminal lysine cannot form cation-pi packing. Therefore, their binding affinities are lower than CIC and LRRK1.

**Figure 4.**
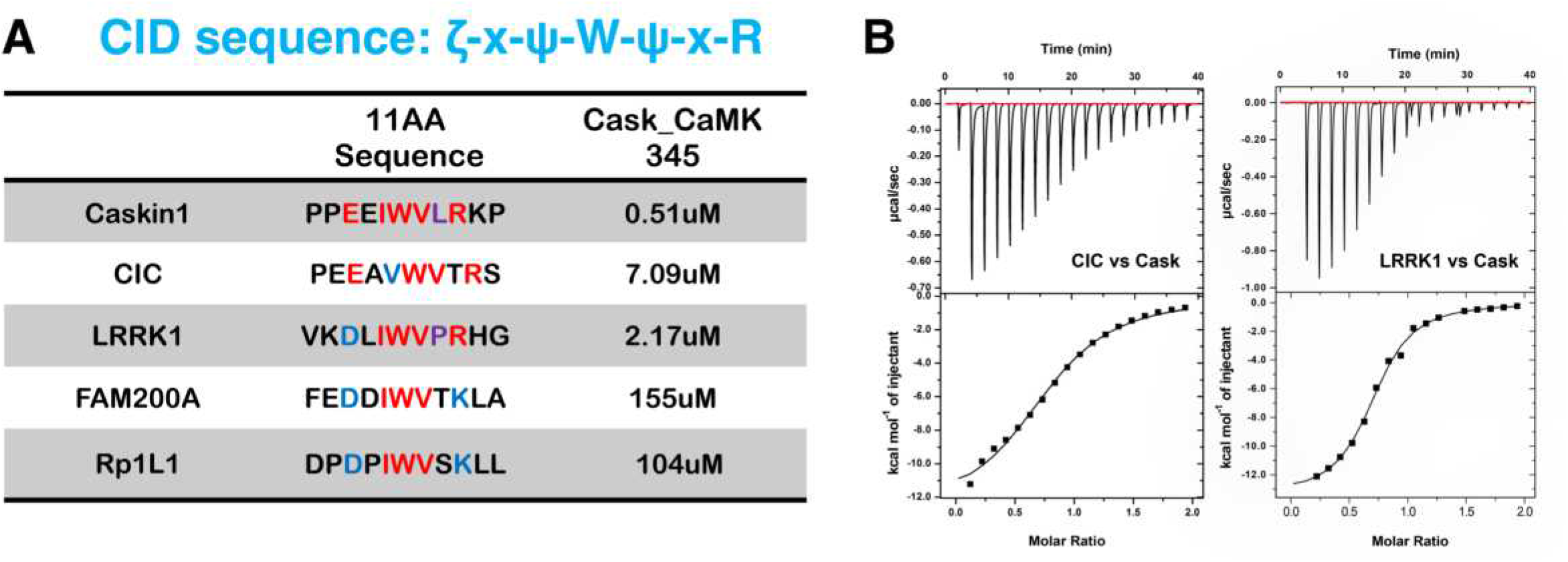
Discovery of the potential new CASK binding proteins and the sequence alignment of their CID sequences According to the CID binding sequence we generated, we identified four potential candidates for CASK_CAMK. ITC-based measurements confirmed the interactions between CID-containing protein and CASK_CAMK.

## Discussion

CASK, as a critical synapse scaffold protein, has many binding targets [1, 8, 13, 20, 34, 36]. For its N-terminal CAMK domain interactions, the CID binding core is essential for all targets [33–35, 37]. By the different binding modes, we divided CASK/CID interactions into three categories: 1. CID binding alone (CASKin1 and Tiam1); 2. CID coupled with helical binding (Mint1) 3. CID coupled with SAM domains binding (LiprinA2) (Figure 5).

**Figure 5.**
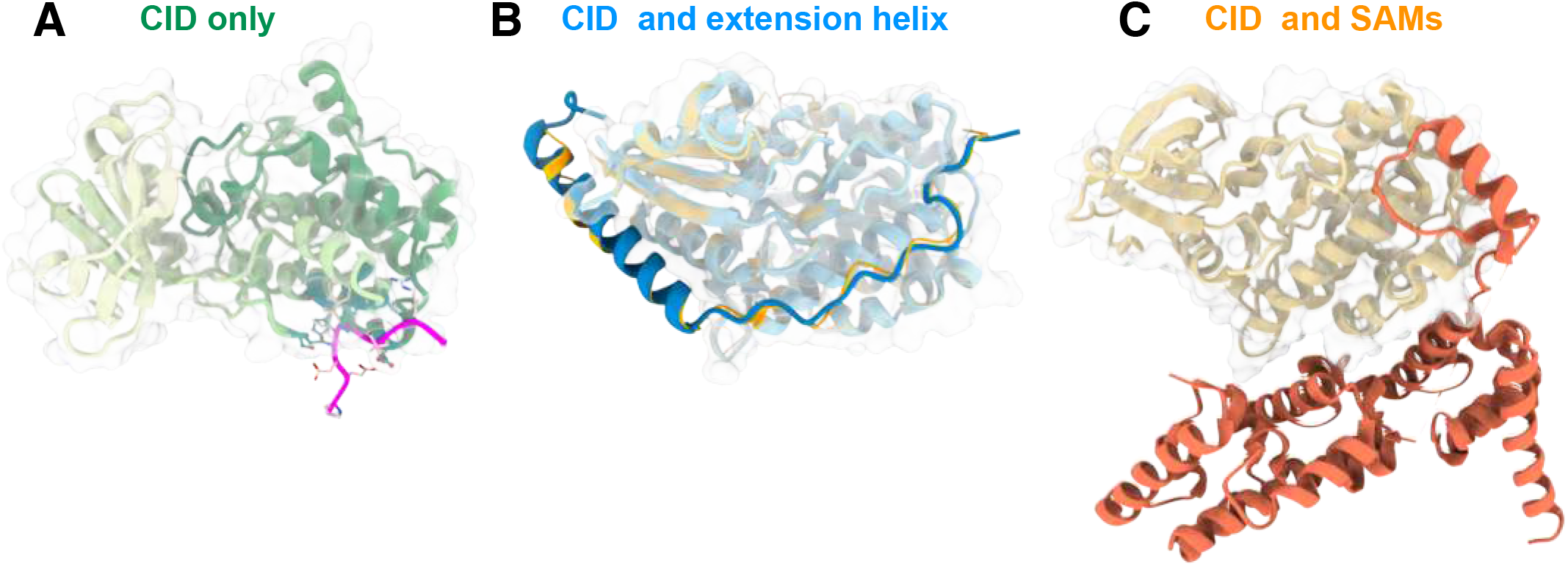
Three models for CASK_CAMK/CID binding.

In this study, we performed structural and biochemical analysis to delight the interaction between Caskin1 and CASK. With the high-resolution crystal structure of the Caskin1_CID/CASK_CAMK complex, we revealed the interaction in detail. A more precise CID sequence has been summarized and two new CASK_CAMK binding partners have been discovered.

Although CASK is expressed in all tissues, the expression levels of CASK are significantly higher in neuron systems than in others [2, 38]. CASK is expressed throughout brain development, with higher protein levels at embryonic stages [39]. The temporal and spatial expression profiles changes let CASK accommodate various targets. On the other hand, CASK enrolls a variety of interaction proteins through its multiple domain organization. These all benefit CASK as a protein platform to assemble numerous targets. Deletion of CASK is lethal and could lower other protein expressions or disrupt the protein distributions.

Caskin1 has a homolog protein Caskin2 [21]. Although the overall domain organization is the same as Caskin1, Caskin2 did not contain the CID sequence. Without the CID sequence, Caskin2 does not bind to CASK and represents a distinguishing function to Caskin1. Previous studies have shown that the Caskin1 SAMs domain is also distinct from Caskin2. Caskin1 SAMs are self-associated into helical polymers[23], while Caskin2 SAMs form high order oligomers repeated in dimers [40].

CIC is a transcriptional repressor, which plays a role in the development of the central nervous system (CNS). LRRK1 is the homolog of LRRK2 [41], which has a complementary expression pattern with LRRK2 in the brain. LRRK2 has the most commonly mutated gene in familial Parkinson’s disease [42]. Comparative low-resolution cryo-EM analysis shows that LRRK1 and LRRK2 could form homodimers possessing similar overall shapes. The same domain organization and similar structure indicate that LRRK1 may have comparable functions with LRRK2. Given this background and the sequence similarity, the two proteins may interact with CASK to play some roles in neuron function.

## Acknowledgements

We thank the Shanghai Synchrotron Radiation Facility (SSRF) BL17U1 and BL19U1 for X-ray beam time.

This work was supported by National Natural Science Foundation of China (Young scientist program) (No. 81801356 and 81802860), China Postdoctoral Science Foundation (Grant No. 2017M612703), Science foundation of Health, Family Planning commission of Shenzhen Municipality (Grant No. SZBC2018017) to Y.W and X.Y, and supported by National Natural Science Foundation of China (No. 31870746), Shenzhen Basic Research Grants (JCYJ20200109140414636), and Natural Science Foundation of Guangdong Province, China (No. 2021A1515010796) to W. L.

## Reference

1. Biederer, T. and T.C. Sudhof, CASK and protein 4.1 support F-actin nucleation on neurexins. J Biol Chem, 2001. 276(51): p. 47869–76.

2. Hata, Y., S. Butz, and T.C. Sudhof, CASK: a novel dlg/PSD95 homolog with an N-terminal calmodulin-dependent protein kinase domain identified by interaction with neurexins. Journal of Neuroscience, 1996. 16(8): p. 2488–2494.

3. Hsueh, Y.-P., et al., Nuclear translocation and transcription regulation by the membrane-associated guanylate kinase CASK/LIN-2. Nature, 2000. 404(6775): p. 298–302.

4. Hsueh, Y.-P., et al., Direct interaction of CASK/LIN-2 and syndecan heparan sulfate proteoglycan and their overlapping distribution in neuronal synapses. The Journal of cell biology, 1998. 142(1): p. 139–151.

5. Fallon, L., et al., Parkin and CASK/LIN-2 associate via a PDZ-mediated interaction and are co-localized in lipid rafts and postsynaptic densities in brain. J Biol Chem, 2002. 277(1): p. 486–91.

6. Lee, S., et al., A novel and conserved protein-protein interaction domain of mammalian Lin-2/CASK binds and recruits SAP97 to the lateral surface of epithelia. Mol Cell Biol, 2002. 22(6): p. 1778–91.

7. Leonoudakis, D., et al., A multiprotein trafficking complex composed of SAP97, CASK, Veli, and Mint1 is associated with inward rectifier Kir2 potassium channels. J Biol Chem, 2004. 279(18): p. 19051–63.

8. Lehtonen, S., et al., Cell junction-associated proteins IQGAP1, MAGI-2, CASK, spectrins, and alpha-actinin are components of the nephrin multiprotein complex. Proc Natl Acad Sci U S A, 2005. 102(28): p. 9814–9.

9. Hsueh, Y.-P., The role of the MAGUK protein CASK in neural development and synaptic function. Current medicinal chemistry, 2006. 13(16): p. 1915–1927.

10. Zhu, J., Y. Shang, and M. Zhang, Mechanistic basis of MAGUK-organized complexes in synaptic development and signalling. Nature Reviews Neuroscience, 2016. 17(4): p. 209–223.

11. Kaech, S.M., C.W. Whitfield, and S.K. Kim, The LIN-2/LIN-7/LIN-10 complex mediates basolateral membrane localization of the C. elegans EGF receptor LET-23 in vulval epithelial cells. Cell, 1998. 94(6): p. 761–771.

12. Chen, K. and D.E. Featherstone, Pre and postsynaptic roles for Drosophila CASK. Molecular and Cellular Neuroscience, 2011. 48(2): p. 171–182.

13. Jeyifous, O., et al., SAP97 and CASK mediate sorting of NMDA receptors through a previously unknown secretory pathway. Nat Neurosci, 2009. 12(8): p. 1011–9.

14. Biederer, T., et al., SynCAM, a synaptic adhesion molecule that drives synapse assembly. Science, 2002. 297(5586): p. 1525–1531.

15. Samuels, B.A., et al., Cdk5 promotes synaptogenesis by regulating the subcellular distribution of the MAGUK family member CASK. Neuron, 2007. 56(5): p. 823–37.

16. Atasoy, D., et al., Deletion of CASK in mice is lethal and impairs synaptic function. Proc Natl Acad Sci U S A, 2007. 104(7): p. 2525–30.

17. Becker, M., et al., Presynaptic dysfunction in CASK-related neurodevelopmental disorders. Translational psychiatry, 2020. 10(1): p. 1–17.

18. Moog, U., et al., Phenotypic spectrum associated with CASK loss-of-function mutations. J Med Genet, 2011. 48(11): p. 741–51.

19. Najm, J., et al., Mutations of CASK cause an X-linked brain malformation phenotype with microcephaly and hypoplasia of the brainstem and cerebellum. Nat Genet, 2008. 40(9): p. 1065–7.

20. Ahn, S.-Y., et al., Scaffolding proteins DLG1 and CASK cooperate to maintain the nephron progenitor population during kidney development. Journal of the American Society of Nephrology, 2013. 24(7): p. 1127–1138.

21. Tabuchi, K., et al., CASK participates in alternative tripartite complexes in which Mint 1 competes for binding with caskin 1, a novel CASK-binding protein. Journal of Neuroscience, 2002. 22(11): p. 4264–4273.

22. Toke, O., et al., Solution NMR Structure of the SH3 Domain of Human Caskin1 Validates the Lack of a Typical Peptide Binding Groove and Supports a Role in Lipid Mediator Binding. Cells, 2021. 10(1).

23. Stafford, R.L., et al., Tandem SAM domain structure of human Caskin1: a presynaptic, self-assembling scaffold for CASK. Structure, 2011. 19(12): p. 1826–36.

24. Wang, Y., et al., Systematic biochemical characterization of the SAM domains in Eph receptor family from Mus Musculus. Biochem Biophys Res Commun, 2016. 473(4): p. 1281–1287.

25. Schindelin, J., et al., Fiji: an open-source platform for biological-image analysis. Nat Methods, 2012. 9(7): p. 676–82.

26. Otwinowski, Z. and W. Minor, [20] Processing of X-ray diffraction data collected in oscillation mode. Methods in enzymology, 1997. 276: p. 307–326.

27. McCoy, A.J., et al., Phaser crystallographic software. Journal of applied crystallography, 2007. 40(4): p. 658–674.

28. Emsley, P., et al., Features and development of Coot. Acta Crystallographica Section D: Biological Crystallography, 2010. 66(4): p. 486–501.

29. Adams, P.D., et al., PHENIX: a comprehensive Python-based system for macromolecular structure solution. Acta Crystallographica Section D: Biological Crystallography, 2010. 66(2): p. 213–221.

30. Murshudov, G.N., et al., REFMAC5 for the refinement of macromolecular crystal structures. Acta Crystallographica Section D: Biological Crystallography, 2011. 67(4): p. 355–367.

31. Chen, V.B., et al., MolProbity: all-atom structure validation for macromolecular crystallography. Acta Crystallographica Section D: Biological Crystallography, 2010. 66(1): p. 12–21.

32. Goddard, T.D., et al., UCSF ChimeraX: Meeting modern challenges in visualization and analysis. Protein Science, 2018. 27(1): p. 14–25.

33. Stafford, R.L., et al., The molecular basis of the Caskin1 and Mint1 interaction with CASK. J Mol Biol, 2011. 412(1): p. 3–13.

34. Zhang, Z., et al., CASK modulates the assembly and function of the Mint1/Munc18-1 complex to regulate insulin secretion. Cell Discov, 2020. 6(1): p. 92.

35. Wu, X., et al., Structural Basis for the High-Affinity Interaction between CASK and Mint1. Structure, 2020. 28(6): p. 664–673 e3.

36. Zhang, K., et al., CASK, APBA1, and STXBP1 collaborate during insulin secretion. Mol Cell Endocrinol, 2021. 520: p. 111076.

37. Wei, Z., et al., Liprin-mediated large signaling complex organization revealed by the liprin-α/CASK and liprin-α/liprin-β complex structures. Molecular cell, 2011. 43(4): p. 586–598.

38. Cohen, A.R., et al., Human CASK/LIN-2 binds syndecan-2 and protein 4.1 and localizes to the basolateral membrane of epithelial cells. The Journal of cell biology, 1998. 142(1): p. 129–138.

39. Hsueh, Y.-P. and M. Sheng, Regulated expression and subcellular localization of syndecan heparan sulfate proteoglycans and the syndecan-binding protein CASK/LIN-2 during rat brain development. Journal of Neuroscience, 1999. 19(17): p. 7415–7425.

40. Smirnova, E., et al., A new mode of SAM domain mediated oligomerization observed in the CASKIN2 neuronal scaffolding protein. Cell Commun Signal, 2016. 14(1): p. 17.

41. Korr, D., et al., LRRK1 protein kinase activity is stimulated upon binding of GTP to its Roc domain. Cell Signal, 2006. 18(6): p. 910–20.

42. Gilks, W.P., et al., A common LRRK2 mutation in idiopathic Parkinson’s disease. The Lancet, 2005. 365(9457): p. 415–416.

