## Supplemental Figures for "Crystal Structure of Caskin1/CASK complex reveals the molecular basis of the binding specificity of CASK_CAMK domain and its binding partners"

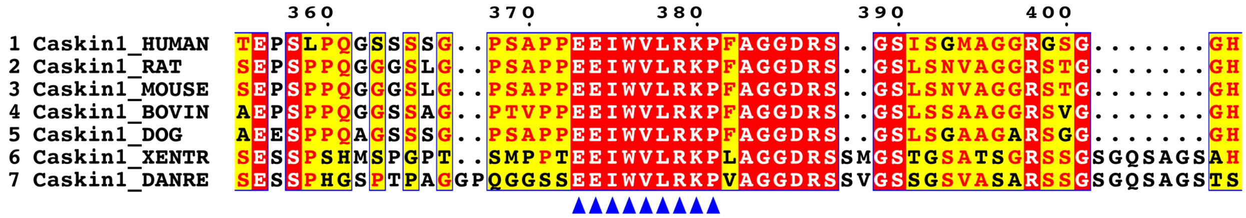


**Supplementary Figure 1.** Sequence alignment of Caskin1 among vertebrate species.

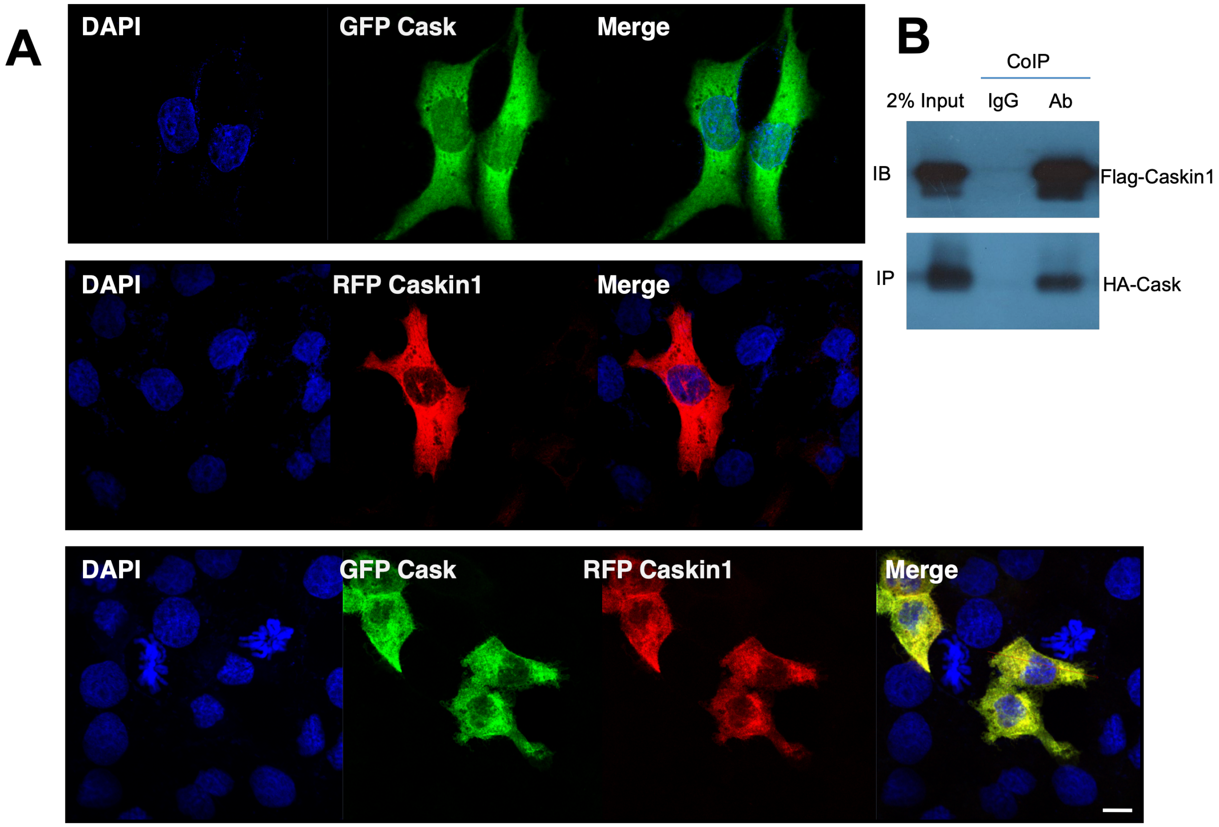


**Supplementary Figure 2.** Caskin1 and CASK co-localized well in HEK293 cells.
A. GFP tagged CASK and RFP tagged Caskin1 well co-localized in the cytoplasm.
B. Co-immunoprecipitation assay showing that the Flag-tagged Caskin1 could be precipitated with HA-tagged CASK.

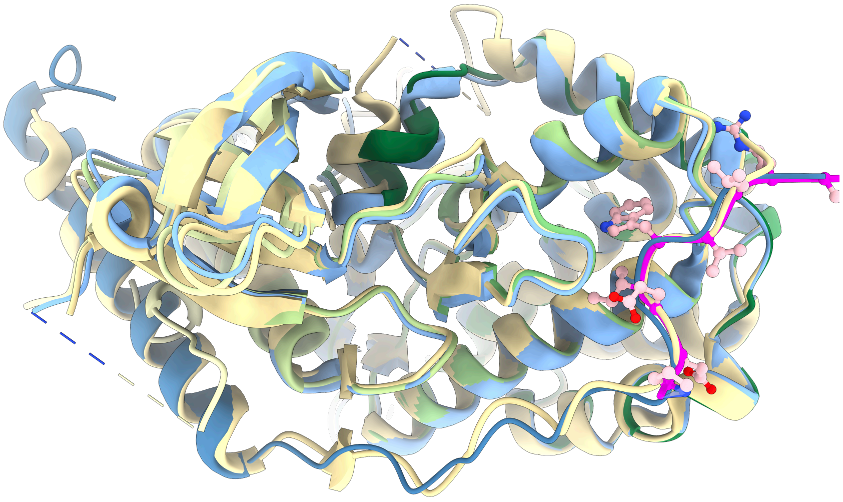

**Supplementary Figure 3.** Superimpose Caskin1/CASK with Mint1/CASK (6LNM, light yellow, and 6KMH, blue) structures. Our model is consistent with extensive structures.
